# Modeling Cognitive Processes with Neural Reinforcement Learning

**DOI:** 10.1101/084111

**Authors:** S.E. Bosch, K. Seeliger, M.A.J. van Gerven

## Abstract

Artificial neural networks (ANNs) have seen renewed interest in the fields of computer science, artificial intelligence and neuroscience. Recent advances in improving the performance of ANNs open up an exciting new avenue for cognitive neuroscience research. Here, we propose that ANNs that learn to solve complex tasks based on reinforcement learning, can serve as a universal computational framework for analyzing the neural and behavioural correlates of cognitive processing. We demonstrate this idea on a challenging probabilistic categorization task, where neural network dynamics are linked to human behavioural and neural data as identical tasks are solved.

## Introduction

An outstanding question in neuroscience is how adaptive behaviour arises out of the continuing interplay between an organism and its surroundings. This interplay is referred to as the perception-action cycle, and is defined by [Fuster, 2004] as “*the circular flow of information from the environment to sensory structures, to motor structures, back again to the environment, to sensory structures, and so on, during the processing of goal-directed behaviour.*” In cognitive neuroscience, various aspects of this perception-action cycle as implemented by the human brain have been examined in isolation, ranging from low-level sensory processes [Wandell and Winawer, 2015] to high-level executive processes [Miller and Cohen, 2001]. Conventionally, investigation of individual cognitive processes boils down to the use of a general linear model to test in what manner a small number of hypothesized task regressors determine the neural activity in particular regions of interest [Friston et al., 1995].

More recently a model-based approach has emerged, where computational models are used to explain the data at hand as subjects are engaged in some cognitive process [Anderson et al., 2008, Ashby and Waldschmidt, 2008, van Gerven, 2016]. In this model-based approach, the researcher puts forward a computational model which serves as an explanandum of neural and behavioural data as a subject engages in a particular task. There are several model families that can provide the backbone of such an explanandum. For instance, hierarchical Bayesian modelling has proven to be very effective in terms of both the derivation of regressors that correlate with neural activity patterns [O’Reilly et al., 2012], as well as in selecting between different models that embody different assumptions [Friston et al., 2003]. The use of Bayesian models is particularly appealing as they also form the basis for normative theories of human brain function [Knill and Pouget, 2004]. However, Bayesian models do not provide insights into how a task might be implemented within the human brain through global interactions between local processing elements.

An alternative approach, which has gained renewed traction in recent years, is afforded by artificial neural network (ANN) models. These models consist of idealized artificial neurons and aim to mimic essential aspects of information processing in biological neural networks. They were originally conceived of as an approach to model mental or behavioral phenomena [Elman et al., 1996, Ritter and Kohonen, 1989]. This study is also referred to as *connectionism* [Hebb, 1949] and was popularized in the 1980’s under the name *parallel distributed processing* [McClelland and Rumelhart, 1989].

Though ANNs were inspired by their biological counterparts [Fukushima, 1980], they have been largely ignored by the neuroscience community in the past few decades. However, due to recent breakthroughs in brain-inspired artificial intelligence, cognitive neuroscientists are now rediscovering their use in furthering our understanding of neural information processing in the human brain. For example, it has been found that the internal states of convolutional neural networks that were trained to categorize objects, are highly informative of neural responses to sensory stimuli in the visual ventral stream [Yamins et al., 2014, Khaligh-Razavi and Kriegeskorte, 2014, Güçlü and van Gerven, 2015]. In this sense, these deep neural networks can be regarded as computational models of the visual ventral stream.

In the present work, we argue that contemporary ANNs are ideal vehicles for modelling increasingly complex cognitive processes that require closing of the perception-action cycle. In this setting, an ANN is taken to represent the silicon brain of an artificial agent that is able to solve the same task as the one being executed by a human subject. That is, the ANN should learn to integrate sensory evidence and generate appropriate actions in order to solve the task at hand. The internal states of the resulting ANN can be used to predict behavioural and/or neural responses of a participant as he is engaged in the same cognitive task. This modelling framework is highly general since it will in principle allow solving of a multitude of cognitive tasks, ranging from basic tasks typically used in monkey neurophysiology [Yang and Shadlen, 2007] to the use of highly demanding tasks such as game playing or free exploration in virtual environments [Mathiak and Weber, 2006, Bellmund et al., 2016, Mnih et al., 2015]. The notion of using ANNs as vehicles for solving cognitive tasks that are of interest to cognitive neuroscientists has recently been proposed by various researchers (e.g. [Mante et al., 2013, Song et al., 2016a, Rajan et al., 2016, Sussillo et al., 2015, Orhan and Ma, 2016]). Particularly by combining neural networks with reinforcement learning one may potentially solve complex cognitive tasks based on reward signals alone, as shown in [Song et al., 2016b].

We further develop the ANN approach to investigate adaptive behaviour. Using a state-of-the-art actor-critic model, we trained a neural network to solve a perceptual decision-making task, requiring categorization of stimuli based on ambiguous sensory evidence. After training, the ANN was presented with the sensations and actions that resulted from a human participant solving the same task. As a proof of principle, we set out to explore the capability of this paradigm in behavioural and neural data in one case participant. For this subject, we show evidence (1) that the ANN performs analogous to the participant in terms of behaviour and (2) that ANN states can be used to predict neural responses as the participant solves the task. Whereas previous work has shown that the internal states of ANNs that were trained to solve a cognitively challenging task show qualitative correspondences with behavioural and neural responses of biological agents [Song et al., 2016a, Song et al., 2016b], here we propose an approach which explains measured behavioural and neural responses for tasks that require cognitive control directly from an ANN’s internal states.

## Human and artificial subjects

### Operationalizing the perception-action cycle

To investigate the perception-action cycle, we designed an experiment in which a (biological or artificial) agent 𝒜 interacts with an environment ℰ. The agent receives observations **x**_*t*_ from the environment and, as a result of changes in ‘brain’ states **z**_*t*_, generates actions **a**_*t*_ that influence the environment. To solve the task, the agent should learn which actions to take at each point in time in order to reach its objectives. It is assumed that learning is driven by reward signals *r*_*t*_, which the agent receives from the environment (see Figure 1 for an overview).

**Figure 1:**
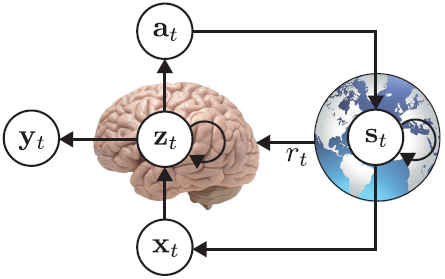
Perception-action cycle. Sensations **x**_*t*_ influence the internal state **z**_*t*_ of an agent 𝒜, resulting in actions **a**_*t*_ (i.e. behaviour) updating the state **s**_*t*_ of the environment ℰ. The agent learns from reward signals *r*_*t*_ that are produced by the environment in response to the agent’s actions. The internal state of the agent can be indirectly sampled using measurements **y**_*t*_.

Let *π*(*a* | **x**) denote a *policy*, which is a function that maps sensory states **x** to discrete actions *a*. That is, we here specialize to the setting where the action can take on one value from a finite set of possible action values. Define the *return*

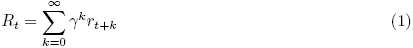

as the total accumulated reward from time step *t* with discount factor *γ* ∈ (0,1] which down-weighs future reward. The agent’s learning goal is to find an optimal policy *π*\* which maximizes the expected return.

In a model-free setting in which the dynamics of the environment are not known, one may resort to reinforcement learning [Sutton and Barto, 1998], where optimal actions are learned solely based on rewards accumulated as an agent explores a task.

We make use of an actor-critic architecture [Sutton and Barto, 1998], which has been linked to reinforcement learning in neurobiology [Barto, 1995, Houk et al., 1995, Joel et al., 2002]. The actor is given by a neural network parametrized by ***θ***, that computes

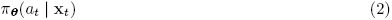

as an approximation of the optimal policy. The critic is given by a neural network parametrized by ***ϕ***, that computes

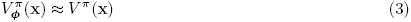

as an approximation of the *value function* ***V***^*π*^ (**x**) = 𝔼[*R*_*t*_ | **x**_*t*_ = **x**], which is the expected return when the current state is **x**. The policy network is used to generate an action given the observations and its parameters are updated in the direction suggested by the critic. Actor-critic models update the policy network parameters ***θ*** by performing gradient ascent on 𝔼[*R*_*t*_]. We refer to the use of neural networks as function approximators for reinforcement learning as *neural reinforcement learning* (NRL).

A complicating factor is that the state of the environment **s**_*t*_ can only be partially observed via the senses through observations **x**_*t*_. This is problematic since, in general, the optimal action may depend on the full state **s**_*t*_ and conditioning on **x**_*t*_ only may result in *perceptual aliasing*, referring to the indistinguishability of different states based on sensations only. One way to solve this issue is by providing a history of observations **x**_*t*_, **x**_*t*−1_,… as input to the neural network. A more elegant solution is to use a recurrent neural network (RNN), whose hidden state depends not only on the input but also on the network’s previous hidden state. Through the use of long short-term memory (LSTM) units [Hochreiter and Schmidhuber, 1997] explicit representations of past events can be kept in memory. Moreover, the RNN can be seen as a representation of brain dynamics that are driven by external sensations and are optimized to maximize expected return. RNNs have been used by [Hausknecht and Stone, 2015] in the context of value-based reinforcement learning.

In the present work, we use asynchronous advantage actor-critic (A3C) learning [Mnih et al., 2016] to train recurrent policy and value networks via gradient updates (see Materials and Methods). After training, the neural network model embodies how a cognitive task can be solved purely through distributed interactions among basic processing elements (i.e. artificial neurons). The model can also be seen as a computational model of how biological agents solve this task in practice. Indeed, in previous work, related models have been shown to agree quite well with behavioural and neuroscientific findings that are known from literature [Song et al., 2016a, Song et al., 2016b]. Our present aim is to show how these models can be conditioned directly on a human participant’s behavioural and neural responses solving the same cognitive task.

### Probabilistic categorization task

Animals continuously make decisions on the basis of their observations of the environment. Learning to map these observations to their corresponding correct decisions is a hallmark of cognition. During a simple decision task, the animal chooses one of two options after evaluating sensory information. The incoming information has to be categorized as belonging to one of two stimulus classes. We here consider a task where this mapping between stimulus class and sensory evidence is probabilistic.

Probabilistic categorization tasks have been widely used to probe learning in human participants. A well-studied paradigm in this literature is the *weather prediction task* [Gluck and Bower, 1988], in which participants are presented with a set of cards depicting abstract shapes, and are asked to decide whether that set of up to four cards predicts one of two outcomes (rain or sunshine). Following each decision, the participant receives feedback, and through this learns the observation-decision mapping. Importantly, all configurations of cards have a certain probability of being paired to rain or sunshine. The weather prediction task has been studied extensively. Work with patients showed that performance on the weather prediction task is primarily dependent on the basal ganglia, and not as much on medial temporal lobe learning system [Knowlton et al., 1994, Knowlton et al., 1996]. Later neuroimaging work confirmed the contribution of the basal ganglia to probabilistic category learning [Shohamy et al., 2004, Poldrack et al., 2001, Ashby and Maddox, 2005, Shohamy et al., 2008, Wheeler et al., 2015, Soltani et al., 2016]. The computational framework of reinforcement learning (RL) has been instrumental in explaining the involvement of the basal ganglia in trial-and-error learning: by learning the future reward value of observations (or *states*) using prediction errors that are conveyed by midbrain dopaminergic neurons [Barto, 1995, Montague et al., 1996, Niv, 2009, Schultz et al., 1997].

Here, we devised a variation on the weather prediction task in which participants are presented with observations sequentially. We consider a task where the ground truth state is either *s*_*t*_ = 0 or *s*_*t*_ = 1, chosen at the start of each trial with probability *p*(*s*_*t*_). At the start of each trial the ground truth state is chosen at random with probability *p*(*s*_*t*_) = 0.5. At each moment in time, the subject has the option to pick state 0, pick state 1 or to wait such as to accumulate another piece of evidence, given by *x*_*t*_. Hence, there are three possible actions to select. Choosing one of the states is realized via button presses whereas refraining from a response for *N* ms is equivalent to a decision to wait.

The evidence is determined by a conditional probability distribution *p*(*x*_*t*_ | *s*_*t*_) which defines the occurrence probability of a piece of evidence given the binary state *s*_*t*_. The evidence variable *x*_*t*_ is also discrete and can take on one of four possible values. We specify the dependence of evidence on state in terms of an odds ratio such that

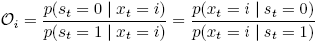

are the odds for the *i*-th piece of evidence under the assumption that *p*(*s*_*t*_) = 0.5. Here, we choose an odds ratio of [0.25,0.75,1.5, 2.5] for each of the four pieces of evidence.

As a result, the symbols each provide partial evidence for either one of the ground truth states. Since the evidence is ambiguous due to its probabilistic nature, evidence needs to be integrated over time in order to infer about the correct ground truth state.

If the subject gives a correct response about the state, this is rewarded with a positive reward (*r*_*t*_ = +15), whereas an incorrect response is punished with a negative reward (*r*_*t*_ = −100). By waiting, the participant can accumulate evidence to decide on the appropriate response. However, waiting for a new piece of evidence also incurs a small negative reward (*r*_*t*_ = −1), so participants are implicitly encouraged to respond fast. The participant is instructed to maximize the number of points scored in each run, and thus should learn the underlying distributions mapping the observations to actions. An overview of this task is given in Figure 2A.

**Figure 2:**
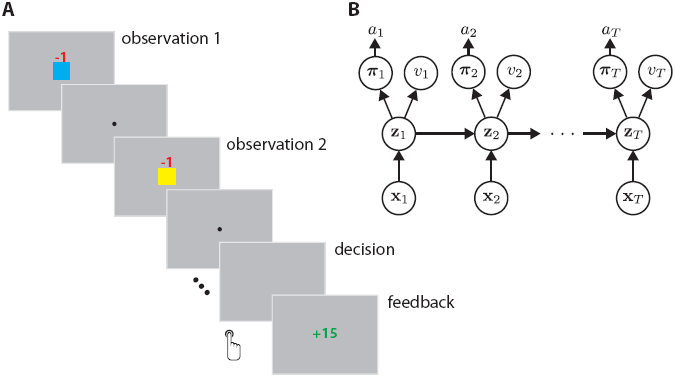
Probabilistic categorization task. **A)** Task design showing the sequential accumulation of evidence encoded by different colour symbols until the agent guesses the underlying state A or B. Red and green numbers indicate negative and positive reward gained by the agent, respectively. **B)** Actor-critic recurrent neural network used to solve the probabilistic categorization task. The states **z**_*t*_ are given by long short-term memory units that keep track of long-term dependencies.

## Results

### Analysis of neural network behaviour

First, we considered to what extent the probabilistic categorization task could be solved using our NRL framework. We trained neural networks consisting of different numbers of hidden units *H* to solve the task (see Figure 2B). We investigated how total reward was related to a range of different number of hidden units *H* (see Figure 3A). At *H* = 45 we observed an optimum, and chose networks with this *H* for the remainder of this demonstration of the framework. Figure 3B shows the distribution of trial lengths for a trained network. This distribution reveals that the network can integrate over more or fewer symbols depending on the uncertainty of the underlying network state.

**Figure 3:**
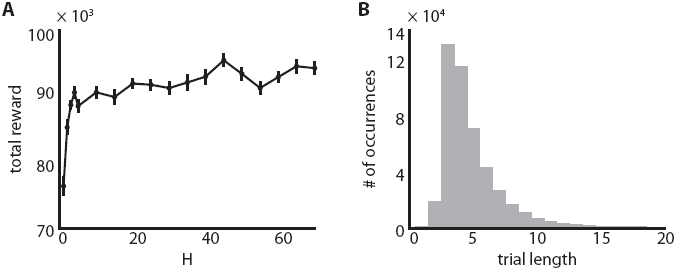
Cumulative reward during task learning. **A)** Total reward after 50.000 time steps as a function of number of hidden units. For each H this presents the average over 42 models trained for 100.000 training steps. Tested on a set of 42 random state and evidence sequences. **B)** Distribution over trial lengths for the best performing network *H* = 45.

To test whether the neural network indeed represents state uncertainty, we correlated state uncertainty given by *p*(*s*_*t*_ = 1 | **x**_*t*_, **x**_*t*−1_,…) with the activations of neural network units. Figure 4 indeed shows that many of the units are positively or negatively correlated with state uncertainty, indicating that an explicit representation of uncertainty is maintained upon which evidence integration may operate. This is in line with recent results which show that rate-based neural networks can maintain explicit representations of probabilistic knowledge [Orhan and Ma, 2016].

**Figure 4:**
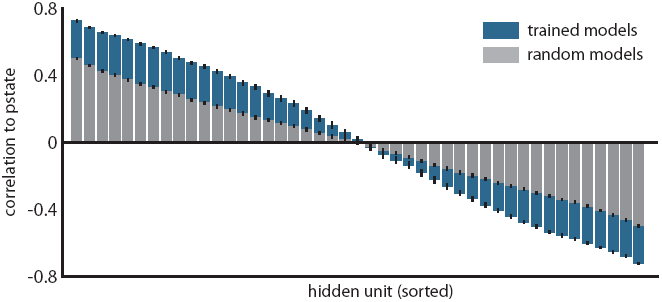
Representation of state uncertainty by activations of neural networks units. Hidden neural network units for trained and random models, sorted by their correlation to state uncertainty *p*(*s*_*t*_ = 1 | **x**_*t*_, **x**_*t*−1_, …). Average over 42 trained and random models.

### Behavioural data analysis

Next, we investigated how well our neural network could account for behavioural data acquired as a human participant solved the same task.

We found that the shape of the human learning trajectory shows a qualitative match to the shape of the learning trajectory of the neural network (see Figure 5A and B): in the initial phase cumulative reward decreases rapidly, until both network and participant learn about the optimal policy, after which cumulative reward starts to increase. The rapid decrease in cumulative or total reward seems to subside around the same time for the subject and the neural network. However, the neural network needs significantly more trials to arrive at a rewarding policy.

**Figure 5:**
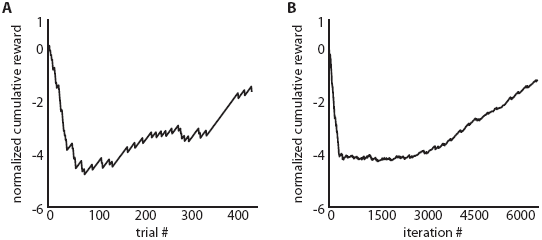
Learning trajectories for a human and an artificial subject. Development of total reward throughout one experiment. Both the human **A)** and the artificial **B)** participant initially have no success, followed by an experimental phase, and finally a near optimal strategy.

Next, we examined to what extent a trained neural network agrees with a subject’s learned policy. This was realised by forcing the model to take the same actions as the subject, while presenting it with the same observations. Let 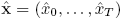 and *â* = (*â*_0_,…, *â*_*T*_) denote the received sensations and performed actions of the subject throughout the experiment. To quantify how well the trained RNN predicts the subject’s behavioural responses (or chosen actions), we use the log probability of the chosen actions:

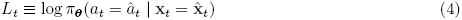

computed at each time step *t*. We compute *L*_*t*_ at each time step by running the model while presenting the same sensory input 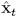, computing log 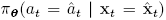, and subsequently clamping the action to the action *â*_*t*_ selected by the subject. If the RNN accurately models subject behaviour, *L*_*t*_ should be larger for the fully trained model compared to a random model.

Initially, the trained model evaluates most subject actions as not beneficial, leading to very low scores (see Figure 6). The policy of the random network, in contrast, is more uniform than that of the trained network, leading to a smaller impact on *L*_*t*_ of wrong decisions made by the participant. In this initial, explorative, stage of the experiment, the participant has almost no information about the correct strategy and the correct choice of actions in each situation. This leads him to act rather randomly and thus acting more similarly to untrained (random) models. In the final stages of the experiment, the participant has found a successful strategy. The trained models agree with participant behaviour much better than random models in this phase.

**Figure 6:**
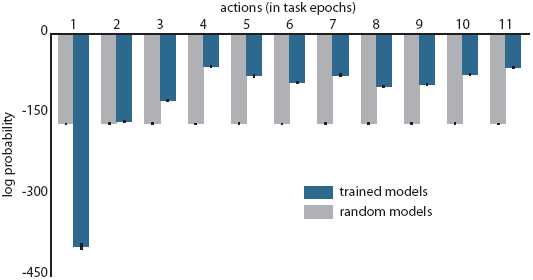
Network evaluation of participant actions. Sums over the log-probabilities of all actions chosen by the participant within epochs of 40 trials. Action log-probabilities are closer to 0 if the ANN model evaluates them as helpful (gathering evidence) or correct (choosing A or B correctly). Results are averaged over 42 trained and 42 random models.

### Neural data analysis

Given the correspondence between model and human behaviour, we asked whether the model could explain variance in the neural data **y**_*t*_ acquired during the task. Specifically, we looked at the post-feedback phase of each trial, during which the participant updated their beliefs about the task and their strategy. To predict the neural signature of reward processing, we set up an encoding model with a number of regressors derived from the network’s hidden states. We used the LSTM hidden states, the entropy, the log-probability of the action chosen by the participant, and the value. We also added a constant to capture offset. These feature regressors were used as a representation of the internal states of the participant during the task, and regressed onto specific post-feedback time bins in the MEG time series using regularized linear regression.

We performed this analysis using feature regressors extracted from random and trained models. We observed that our encoding models were predictive of a post-feedback signal in the frontomedial region, peaking within the 280 ms − 320 ms time window at three adjacent sensors (see Figure 7). The features extracted from trained models predicted neural responses better than those from random models. While the overall effect size is low, it consistently explains variance of the post-feedback signal for trained models. This indicates that neural network features can be used to predict neural signals during a cognitive task.

**Figure 7:**
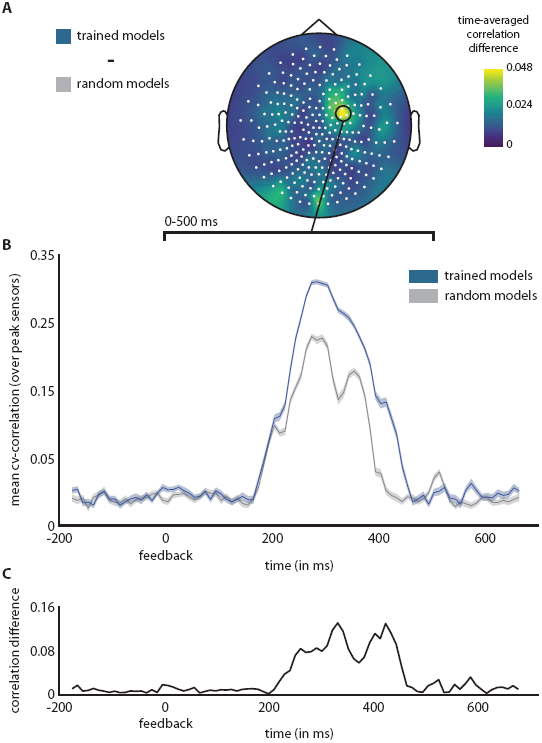
Correlations between predicted and measured MEG activity at feedback presentation. **A)** Spatial representation of correlation difference between trained and random models, time-averaged over the visual presentation of feedback (0–500 ms). **B)** Time-resolved correlation traces for trained and random models at the peak sensors in A). Averaged results are shown for the 42 trained and random models. **C)** Time-resolved correlation difference between trained and random models in B).

## Discussion

In the present work, we describe a neural modelling framework that can be used to study the separate components of the perception-action cycle. Here, we demonstrate this framework with a probabilistic categorization task. We show that artificial neural networks can be trained to solve this challenging cognitive task. Importantly, the network states can be used to explain behavioural and neural responses from human participants solving the same task. We argue that our framework can provide new insights into cognitive processes by relating network representations to human representations of an experimental task.

When comparing learning trajectories, we found that the ANN and the human subject showed similar learning trajectories. Note, however, that the human subject required less time to find a working solution that enabled him to accumulate reward. This may be explained in part by the fact that our case subject was not completely naive: unlike the neural network he did not have to learn the meaning of the buttons (of the three possible actions), and was informed that he had to use the evidence in order to make a decision. In addition he could rely on human a *priori* knowledge [Lake et al., 2016]. In a pilot in which we asked participants to learn the task without instructions (data not shown), only presenting them with the goal of maximizing the score, we observed large inter-subject variability, with some subjects learning the task efficiently, some subjects using clever strategies to learn about the nature of the task (e.g. asking for very long symbol sequences) and some subjects not learning the task at all. While it is naive to assume that the ANN employs the same strategy to learn the task, the learning trajectories themselves can be used to select among different learning schemes and neural network architectures when the objective is to accurately model human behaviour. Moreover, optimizing the correspondence in learning trajectories by varying for example the learning rate or *β* parameter, which modulates the tradeoff between exploration and exploitation, may shed new light on individual differences in learning strategy.

In our analysis of the behaviour of ANNs that solve the probabilistic categorization task, we found that ANN internal states code for the uncertainty about the state of the environment. Such coding is essential, as it allows the ANN to integrate evidence such as to arrive at an optimal decision. Previous work has shown that rate-based neural networks are indeed able to explicitly represent uncertainty (i.e. graded beliefs) in order to optimally solve a task [Shen and Ma, 2016, Rao, 2004]. As such, ANNs can provide an account at the implementational level of how normative theories derived from Bayesian principles can be realized in the human brain.

We also demonstrated how ANNs trained to solve cognitively challenging tasks can be used to predict neural responses. This expands on previous work in sensory neuroscience, where deep neural networks have been used to predict neural responses to naturalistic sensory input [Güçlü and van Gerven, 2015]. The present work shows how the same can be achieved in decision neuroscience, using neural network regressors to predict neural responses in tasks that require cognitive control. We stress that the goal of the present work is limited to providing an illustration of how neural networks can be used to study neural correlates of cognitive control in principle. The use of one subject as a case study together with the relatively low effect size precludes a detailed analysis of our present findings. Still, it is interesting to see that the network features seem to predict an ERF component elicited by the feedback signal. This component may be linked to prior studies showing that post-stimulus P3a potentials are related to the magnitude of belief updates in reinforcement learning settings [Bennett et al., 2015] rather than the magnitude of presented feedback. An important caveat is that both trained and random models are predictive of the ERF component. This could simply reflect prediction of the ERF component by the constant term used in the forward model. However, the trained model does explain more variance than the random model. Whether or not this points towards a direct link between ANN regressors and neural response variability, remains an open question at present. To fully exploit the potential of our framework to probe neural correlates of decision-making, further research that goes beyond the illustration provided by this case study is needed.

A challenge for future research is to come up with learning schemes and neural network architectures that combine human-like performance levels with biological plausibility. In terms of network architectures, we have used a particularly straightforward implementation of a recurrent neural network architecture. Such architectures may be made more realistic by for example allowing high-dimensional sensory input by using convolutional layers, allowing multiple states of processing as in deep reinforcement learning [Mnih et al., 2015, Lillicrap et al., 2015], as well as adding biophysical constraints that take properties of human sensory or motor systems into account (see e.g. [Lillicrap and Scott, 2013, Sussillo et al., 2015]). Furthermore, the range of possible cognitive tasks which can be tackled by neural reinforcement learning approaches can be extended by using continuous or hybrid policies [Lillicrap et al., 2015]. In this way, we can model not only discrete decisions but also continuous changes in sensory and/or motor parameters that are required to optimally solve particular tasks, as in active sensing [Yang et al., 2016]. The present actor-critic model shares interesting properties with biological learning. It has been suggested that dorsolateral prefrontal cortex may correspond to the policy network whereas the basal ganglia or orbitofrontal cortex may correspond to the value network [Song et al., 2016a]. New advances in biologically plausible learning algorithms may further bridge the gap between learning in biological and artificial agents [Miconi, 2016, Lillicrap et al., 2014, Rombouts et al., 2015, Scellier and Bengio, 2016].

Reflecting on past and present results, it is exciting to observe that neural networks are being re-appreciated as models of human cognitive processing. New advances in ANN research make it possible to train neural networks to solve increasingly complex tasks, approaching (and sometimes surpassing) human-level performance. Due to these advances, ANN dynamics can now be linked to behavioural and neural data acquired as human subjects (or animals) perform the same challenging task. This convergence between *artificial intelligence*, providing increasingly sophisticated learning algorithms and architectures, and *neuroscience*, providing biological constraints on such algorithms and architectures, is expected to lead to new insights in cognitive processing in humans. Tasks of interest may range from simple, like the probabilistic categorization task described here, to complex, like challenging video game environments in which many strategies are possible.

## Materials and Methods

In the following, we describe the architecture of our recurrent neural network, the experimental procedure, the MEG data analysis and the structure of the encoding model.

## Neural network

### Network architectures

In the probabilistic categorization task, we make use of an RNN consisting of an input layer, a hidden layer and an output layer. The input layer consists of *K* binary artificial neurons and represents the presented evidence symbol using a one-out-of-*K* coding scheme. The hidden layer consists of *K* long short-term memory (LSTM) units with forget gates that are designed to explicitly store long-term dependencies [Gers, 2001], [Hochreiter and Schmidhuber, 1997]. The LSTM units can be viewed as the component of our network resembling working memory. They learn to store information about the sequence of evidence. The output layer consists of *M* + 1 output neurons with *M* neurons coding for the policy and one neuron coding for the value function. The policy is represented by the *M* neurons using a one-out-of-*M* coding scheme in conjunction with a softmax activation function which maps linear activations to action probabilities. Based on these action probabilities an action *a*_*t*_ is selected. The value function is represented by one neuron with a linear activation function. The complete framework is implemented using the Python framework Chainer^1^.

### Long short-term memory units

An LSTM unit has the ability to remove or add information to the *cell state c_t_*, as regulated by the following gates:

- An *input gate i*_*t*_ = *σ*(**w**_*i*_ · [**h**_*t*−1_, **x**_*t*_] + *b*_*i*_) which decides what information to store.
- A *forget gate f*_*t*_ = *σ*(**w**_*f*_ · [**h**_*t*−1_, **x**_*t*_] + *b*_*f*_) which decides what information to forget.
- An *output gate o*_*t*_ = *σ*(*W*_*o*_ · [**h**_*t*−1_, **x**_*t*_] + *b*_*o*_) which decides what information to pass on.

The *cell input* depends on the hidden states **h**_*t*−1_ and is given by 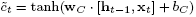. Using the input and forget gates the cell state is updated to 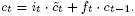. Finally, the *hidden state* of the LSTM unit is computed as *h*_*t*_ = *o*_*t*_ *·* tanh(*c*_*t*_).

### Training procedure

All neural networks were trained using asynchronous advantage actor-critic (A3C) learning [Mnih et al., 2016]. Details of the applied training procedure are described in the following.

Let *J*(***θ***, ***ϕ***) be any policy objective function. Policy gradient algorithms search for a gradient of the policy *π*(*a* | **x**; **θ**) with respect to parameters **θ**:

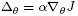

According to the policy gradient theorem, the policy gradient is given by

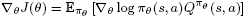

where ∇_*θ*_ log *π*_*θ*_ (*s*, *a*) is the *score function* and 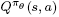 is the *action-value function*.

Let 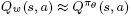 be an estimate of the action-value function. Actor-critic (AC) algorithms consist of a *critic* which updates action-value function parameters *w* and an *actor* which updates policy parameters *θ* in the direction suggested by the critic.

AC algorithms follow an approximate policy gradient

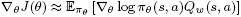

which becomes exact when the compatible function approximation theorem is satisfied (value function approximator is compatible to the policy and value function parameters minimize the mean squared error). By subtracting a baseline function *B*(*s*) from the policy gradient we can reduce the variance of our estimator without changing the expectation. A good baseline function is the value function 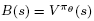 so we can write

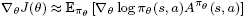

where 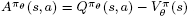 is the *advantage function*. Hence, the critic should really estimate 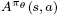. In practice, we use the return accumulated for *n* time steps as an estimate of the action-value function. For a discrete action space, we use a softmax policy:

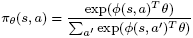

We wish to compute

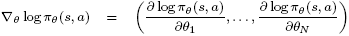

where

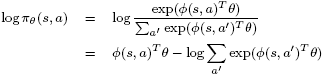

is the log softmax function.

### Gradient updates

The policy is improved by performing (approximate) gradient ascent on the expected return. In practice, the gradient updates are implemented by an *n*-step look-ahead when interacting with the environment. In this case the return is given by

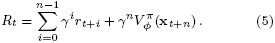

Let 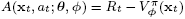 be the estimated *advantage* of performing action *at* when observing **x**_*t*_. Let

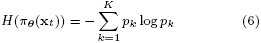

with *p*_*k*_ = *π*_***θ***_(*a*_*t*_ = *k* | **x**_*t*_) be the entropy of the (stochastic) policy. The policy gradient updates are given by:

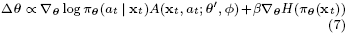

where the contribution of the entropy is controlled by hyperparameter *β* and where log *π*_***θ***_(*a*_*t*_ | **x**_*t*_) is known as the *score function.* The gradient updates push the parameters towards or away from the action *a*_*t*_ depending on the sign of the advantage. The entropy term aids exploration by preventing convergence towards a single action [Williams and Peng, 1991].

Note that the policy updates rely on the current estimate of the value function through the dependence on the advantage. The value function 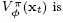 is improved by performing (approximate) gradient descent on the squared advantage using the following update of the parameters ***ϕ***:

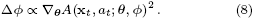

We refer to this squared advantage as the *surprise*, since it quantifies the discrepancy between the reward obtained when performing an action in a particular states and the value of that state according to the model. Models were trained using backpropagation through time as implemented in Chainer. To perform the gradient updates we used RMSProp as an optimizer (learning rate of 0.01, gradient normalization term *ε* of 0.1 and momentum decay rate *α* of 0.99). Initial weights of the network were drawn from scaled Gaussian distributions as described in [He et al., 2015]. To facilitate convergence during model training, we made use of an asynchronous implementation where gradient updates were run in parallel on sixteen CPU cores with all cores applying parameter updates to a common model [Mnih et al., 2016], however learning them in individual environments. Such asynchronous updates stabilize online reinforcement learning by bypassing strong correlations between successive updates in a single environment.

### Trained and random models

In this case study, we make the assumption that the human task representation is more similar to an internal representation in a fully trained neural network than to a representation from a random one. Behavioural and encoding results are analyzed by comparing between models with fully trained and random weights.

Since training of neural network models has random components, one single model can not be considered representative. We thus summarize the encoding and behavioural results over 42 trained and 42 random models, all having an identical architecture with *H* = 45 hidden units. Each of the fully trained models underwent 100,000 training iterations, leading to near-perfect performance in the probabilistic categorization task. In contrast, random models did not undergo any optimization of their randomly set initial weights.

### Physiological correlates

Various descriptors extracted from the RNN can be regarded as the current representation of a state in a given trial. Below, we formally outline the descriptors we later use as a representation in the encoding framework.

### Entropy

The entropy of the stochastic policy is a measure of uncertainty of the optimal action to perform in the current state according to the actor. At each point in time, the entropy is computed as:

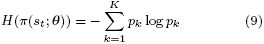

with *p*_*k*_ = *π*(*a*_*t*_ = *k* | **s**_*t*_ = **ŝ**_*t*_; ***θ***).

### Log-probability of the chosen action

The probability of the subject’s decision in the network’s policy, given the current problem or evidence sequence. This can be regarded as a measure for how much the neural network agrees with the subject’s decision:

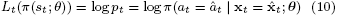

### Value

The value according to the critic is a measure of the expected return. At each point in time, the value is given by:

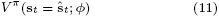

### LSTM states

The current states of the LSTM units, corresponding to the hidden layer. They learned to store relevant information about the previous evidence sequence and can be understood as an internal representation of the current problem.

## Experimental procedure

### Participant

One participant (male, age 30) with normal vision participated in the study and gave written informed consent in accordance with the Declaration of Helsinki. The study was approved by the local ethical review board (CMO region Arnhem-Nijmegen, The Netherlands) and was carried out in accordance with the approved guidelines.

### Stimuli

The stimuli were generated using MATLAB (version 2016a) and the Psychophysics Toolbox [Brainard, 1997]. Stimuli were displayed using an LCD projector (1024 × 768 resolution, less than 1ms presentation delay) against a uniform grey background. Visual stimuli consisted of centrally presented coloured squares (the evidence symbols, 1^°^), rewards (font size 26) and a central fixation point (radius, 0.075^°^).

### Task instructions

The learning phase of the participant and that of our neural network were kept as similar as possible by instructing our participant minimally about the nature of the task. Our participant was instructed that his main goal was to maximize his score. We informed him that there were two answering buttons, and that evidence would consist of four different colours appearing automatically if he decided to wait. The information about four different observations corresponds to our ANN receiving input as a four-element vector that presents a one-hot encoding for the symbols. The participant was prepared for the temporal structure of a trial (such as inter-trial intervals and blinking periods) prior to the experiment by showing him two example trials that were non-informative about the actual evidence distribution. The subject was instructed that he would receive an additional monetary reward if he scored above 0 points within a run. He maintained fixation on the central fixation point throughout all runs.

### Probabilistic categorization task

Our participant performed twenty-four separate runs (of 20 trials each) of a probabilistic categorization task. Due to technical problems, the data of the first two runs were discarded. During the task, each trial started with the onset of the fixation point (800 ms). Subsequently, an evidence symbol was presented centrally for 1.75 s, with a negative reward cue (−1) directly above it (presented for 500 ms). Following the symbol presentation, there was a variable inter-stimulus interval (ISI, between 0.5–1 s) before the next symbol was presented. In total, a trial could consist of 40 sequentially presented symbols. The participant was instructed to press one of two buttons to indicate their decision on the basis of the presented visual stimuli. This response was possible at any moment during the presentation periods and the ISIs between them. Upon a button press, there was a variable interval (between 1.5–2 s), after which the trial feedback was presented centrally for 500 ms. This feedback consisted of a number indicating the score obtained on that trial. This score was calculated by adding the response reward (+15 for a correct response, -100 for an incorrect response) to the number of presented symbols in that trial (-1 for every symbol). The trial score was presented in green if positive, and in red if negative. If no response was given during a trial, this was regarded as an incorrect response. After feedback presentation, a blank screen intertrial interval (ITI) was presented (randomly jittered between 1.5–3.5 s), during which participants could blink.

### MEG data acquisition

Continuous whole-brain activity was recorded using a CTF 275-channel MEG system (CTF MEG systems, VSM MedTech) at a sampling rate of 1200Hz. 4 channels were malfunctional at the acquisition time and had to be removed from the data, leading to 271 channels. Ear canal and nasion markers were used to continuously monitor head position via a real-time head localizer [Stolk et al., 2013]. When head position deviated >5 mm from the origin position (registered at the beginning of recording), subjects readjusted to the origin position at the next break between runs. An EyeLink 1000 eyetracker (SR Research, Ottawa, Canada) was used to continuously track the left eye to detect eye blinks and saccades.

## MEG data analysis

### MEG preprocessing

All MEG preprocessing steps used functionality and standard data processing pipelines from the Matlab framework FieldTrip^2^ [Oostenveld et al., 2011]. A DFT filter was applied to remove 50 Hz line noise and its harmonics at 100 Hz and 150 Hz, with a data padding of 10 s. Data was demeaned with a baseline taken from -250 ms and -50 ms before every trial. Trials were defined from -200 ms to 700 ms around the visual presentation of reward (+15) or punishment (−100), called *feedback* subsequently.

We downsampled the data to 300 Hz before the subsequent Independent Component Analysis (ICA) to stabilize this step numerically. Before ICA, we inspected trial summary statistics visually to reject those trials with overly high variance and kurtosis, as a first pass for rejecting trials with eye blinks and other muscle movements [Delorme et al., 2007]. For detecting trials with muscle artefacts, data was high-pass filtered at 100 Hz; then summary statistics were inspected. Using this procedure we rejected 30 feedback presentation trials of our case subject. ICA had the most influence on cleaning the data from noise from heart beat, blink, eye movement, and muscle artefacts. Independent components reflecting heart rate, and movements in the eye region were rejected by visual inspection on a trial-by-trial basis. This visual rejection of ICA components was supported by taking the independent components most correlated with EOGh, EOGv and ECG activity into account. After removing the components, we inspected the data visually once more, with the same procedure that we used before running the ICA. The cleaned data was then baseline corrected with a window of -250 ms to -50 ms before the trial, and finally lowpass filtered at 100 Hz.

The outcome of this preprocessing procedure were 410 trials (30 of 440 rejected) at 300 Hz defined around feedback presentations for each of the 271 functioning sensors.

### Extracting sensor responses

For encoding, we described the sensor signal at a temporally dense distance of 10 ms within the 900 ms window defined around the feedback presentation. At each of these time points, we took the subsequent average of the amplitude in a time window of 45 ms. The encoding model attempts to learn predicting these windowed amplitude averages at a specific point in time (*sensor responses*) based on a problem encoding with neural network parameters.

### Problem representation

We extract features from a neural network model by presenting it with the same series of trials as the subject. The neural network (the artificial subject) is clamped to make a decision for the presented problem at the same time as our case subject. Effectively, this means that the artificial subject will request and be presented with the same evidence symbols as our human subject before making a decision. The experiment with the artificial subject applies the set of fully trained weights and does not involve further training, so the network is not informed about actual decisions or rewards. The artificial subject’s internal states are influenced solely by the presented evidence. Each new piece of evidence changes the artificial subject’s internal representation of the current problem, its rating of the available actions and its expected value after making a choice. The features used as an encoding - as the *representation* of the current problem - are these internal neural network states and expectations at the time point of a decision, given the previous evidence sequence. We associate them with sensor responses at specific time points using the encoding model approach [Naselaris et al., 2011]. We extracted the following *M* = 49 regressors, formally defined in section *Physiological Correlates*, from a given neural network with our optimum of *H* = 45 hidden LSTM units:

- the states of the 45 LSTM units that comprise the RNN,
- the entropy of the current policy given the presented evidence pieces (Equation 9),
- the log probability of the action chosen by the subject (Equation 10) (the artificial subject’s *rating* of this action),
- the expected value (Equation 11),
- a constant 1-vector that serves as a bias catching consistently occurring event-related fields.

Note that we do not use any explicit information about the observed reward other than the current internal expectations of the neural network. The above representation was extracted for each trial, and for each of the 42 trained and random models. Encoding results are summarized over these sets of models, and compared between fully trained and random.

### Regression model

We trained a series of independent linear ridge regression models, one for each sensor and time point surrounding feedback presentation. Each of these linear models maps a given set of features to a sensor response at a particular time point. A ridge regularizer was used to avoid overfitting.

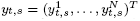 be a vector containing responses for every trial, for one specific sensor s at one specific time bin *t*. Let Φ = (Φ^1^,…, Φ^*N*^)^*T*^ be a *N* × *M* matrix representing the feature vectors for each trial. Our linear model’s regression coefficients 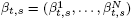 are then estimated by:

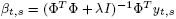

where *λ* ≥ 0 is the regularization strength.

The performance of a model at a specific sensor and time point is the Pearson’s correlation between the *measured* and the *predicted sensor response*, predicted by the model with optimal λ. This performance is estimated on the held-out folds in each of 5 top-level cross validation iterations. Folds were selected randomly from all trials. The final correlation for each sensor-time pair is the average over the held-out fold test correlations, corrected with Fisher’s z-transformation. Nested within each of these 5 cross-validation runs was one further cross validation that was used for selecting the regularization strength *λ* (best *λ* in *k* = 4 folds).

For every trained and random neural network model, this process lead to one temporal series of prediction-activity correlations for each of the 271 sensors, at 85 time bins defined within a -200 ms to 700 ms window around feedback presentation.

## Footnotes

1 http://chainer.org

2 Version: June 8th 2016, git --short hash ebc7d0b. See http://www.fieldtriptoolbox.org.

